# Mu and Delta Opioid Receptors Modulate Inhibition within the Prefrontal Cortex Through Dissociable Cellular and Molecular Mechanisms

**DOI:** 10.1101/2024.10.17.618870

**Authors:** Rebecca H. Cole, Max E. Joffe

## Abstract

Aberrant signaling within cortical inhibitory microcircuits has been identified as a common signature of neuropsychiatric disorders. Interneuron (IN) activity is precisely regulated by neuromodulatory systems that evoke widespread changes in synaptic transmission and principal cell output. Cortical interneurons express high levels of Mu and Delta opioid receptors (MOR and DOR), positioning opioid signaling as a critical regulator of inhibitory transmission. However, we lack a complete understanding of how MOR and DOR regulate prefrontal cortex (PFC) microcircuitry. Here, we combine whole-cell patch-clamp electrophysiology, optogenetics, and viral tools to provide an extensive characterization MOR and DOR regulation of inhibitory transmission. We show that DOR activation is more effective at suppressing spontaneous inhibitory transmission in the prelimbic PFC, while MOR causes a greater acute suppression of electrically-evoked GABA release. Cell type-specific optogenetics revealed that MOR and DOR differentially regulate inhibitory transmission from parvalbumin, somatostatin, cholecystokinin, and vasoactive intestinal peptide-expressing INs. Finally, we demonstrate that DOR regulates inhibitory transmission through pre- and postsynaptic modifications to IN physiology, whereas MOR function is predominantly observed in somato-dendritic or presynaptic compartments depending on cell type.

**Significance Statement:** The endogenous opioid system regulates behaviors that rely on prefrontal cortex (PFC) function. Previous studies have described Mu and Delta opioid receptor expression within cortical GABAergic interneurons, but a detailed understanding of how opioids regulate different interneuron subtypes and cortical microcircuits has not been reported. We use whole-cell patch-clamp electrophysiology, genetically engineered mice, and optogenetics to assess MOR and DOR regulation of PFC inhibitory transmission, demonstrating that MOR and DOR inhibition of interneurons display qualitative and quantitative variation across GABAergic circuits within mouse prelimbic PFC.

## Introduction

The prefrontal cortex (PFC) is critical for regulating reward, motivation, and affective processing—dysfunction of which represent core features of neuropsychiatric disorders. PFC function is maintained by a diverse pool of GABAergic interneurons (INs) that control pyramidal cell spike-timing, synaptic plasticity, and oscillatory states [76]. Disruptions to PFC INs have been implicated in the etiology of schizophrenia, major depressive disorder, and autism, among other diseases [4, 5, 21, 25, 28, 31, 39, 48, 58]. Conversely, manipulations that target PFC INs represent attractive strategies for the treatment of psychiatric symptoms, motivating a better understanding of the biological processes and signaling elements that regulate these circuits.

Neuropeptides and their cognate G protein-coupled receptors (GPCRs) are preferentially expressed within cortical INs. GPCR modulation of INs has been well-documented [8, 9, 27, 30, 42, 43, 68] and can modify behaviors relevant to psychiatric disease [24, 37, 54]. The endogenous opioid peptides and receptors comprise perhaps the most prominent neuropeptide system within the forebrain. Opioid signaling is typically transmitted through Gα_i/o_-dependent signaling that regulates pre- and postsynaptic functions to suppress neuronal activity [26, 64]. The opioid system plays a major role in regulating physiological processes related to cognition, sensation, and behavior.

Aberrant opioid signaling has been proposed in biological hypotheses related to several diseases that affect PFC function [10, 57, 62, 66, 77]. Furthermore, medications that engage opioid receptors are used to treat addictive disorders and pain, with experimental opioid-based treatments under investigation for major depressive disorder and other diseases [1, 11, 13]. Despite these compelling associations, we have a limited underlying of the mechanisms through which opioids regulate PFC IN function. The Mu opioid receptor (MOR) and Delta opioid receptor (DOR) display predominant expression within INs, and only weak expression in excitatory pyramidal cells [69]. A handful of studies spread across frontal cortex subregions have described how MOR or DOR regulate discrete aspects of inhibitory transmission [2, 12, 15, 18, 36, 45]; however, a comprehensive and comparative assessment of MOR and DOR regulation of PFC has not been reported.

In this study, we use whole-cell patch-clamp electrophysiology, slice pharmacology, and cell type-specific optogenetics to characterize MOR and DOR modulation of prelimbic (PL)-PFC inhibitory transmission. We show that MOR and DOR signaling differentially suppress spontaneous inhibitory transmission through different molecular mechanisms. Furthermore, we find that MOR and DOR suppress action potential-dependent GABA transmission through a set of dissociable mechanisms. We next surveyed MOR and DOR expression and functions throughout GABAergic INs, finding that MOR and DOR actions are synapse-specific and show differences in presynaptic versus somato-dendritic function across different IN subtypes. Overall, this study advances our understanding of how Mu and Delta opioid signaling regulate local inhibitory microcircuitry in the mouse PL cortex, providing a framework for understanding synapse-specific perturbations in disease states.

## Materials and Methods

### Mice

Adult (>10 week) male and female genetically engineered mice were bred at Charles River Laboratory (Wilmington, MA) and allowed to acclimate to the animal housing facility for at least one week before experiments. Mice were identified as male or female based only on external genitalia. PV-Cre [33], PV-Cre/SST-Flp [32], CCK-Cre/VGAT-Flp [74], and VIP-Cre/SST-Flp [32] mice were used for optogenetic experiments. PV-Cre-tdTomato, SST-Flp-tdTomato, CCK-Cre/VGAT-Flp, and VIP-Cre-tdTomato mice [33, 40, 53] were used to target IN types for holding current recordings. All mice were bred on a congenic C57BL/6J background. Mice were provided with food and water *ad libitum* and maintained on a 12-hour light/dark cycle. Approximately 75% of experiments were conducted after sacrificing mice during the light phase (lights on at 7:00 am) and 25% conducted during the dark phase (lights off at 10:00 am). No significant effect of phase was observed, and data are pooled across light cycle. All experiments were performed in groups of sex-matched littermates housed 2-5 per cage. Experiments were approved by the University of Pittsburgh IACUC and conducted in accordance with NIH Guidelines.

### Whole-Cell Patch-Clamp Electrophysiology

Whole-cell patch-clamp recordings were performed as previously described [38]. Mice were deeply anesthetized under isoflurane anesthesia and rapidly decapitated. Acute 300-micron coronal slices containing the PL-PFC were prepared in N-methyl-D-glucamine (NMDG)-based cutting solution (in mM: 93 NMDG, 20 HEPES, 2.5 KCl, 0.5 CaCl_2_, 10 MgCl_2_, 1.2 NaH_2_PO_4_, 25 glucose, 5 Na-ascorbate, and 3 Na-pyruvate) at room temperature and immediately transferred to 32°C NMDG solution for 10 minutes. Slices then recovered for 1 hour in artificial cerebrospinal fluid (aCSF) (in mM:119 NaCl, 2.5 KCl, 2 CaCl_2_, 1 MgCl_2_, 1 NaH_2_PO_4_, 11 glucose, and 26 NaHCO_3_) (20-24°C) before being transferred to a recording chamber and perfused (2 mL/min) with heated (30-32°C) aCSF. All solutions were oxygenated with 95% O_2_/5% CO_2_. Data were acquired using a Multiclamp 700B amplifier and pClamp11 software (Molecular Devices). Putative pyramidal neurons were identified based on characteristic membrane properties (i.e. high capacitance, low input resistance, spike-firing adaptation) and INs were identified by tdTomato or GFP fluorescence. All recordings were performed in PL-PFC L2/3. Cells were patched with a borosilicate glass micropipette pulled to 3-6 MΩ with a horizontal electrode puller (P-1000, Stutter Instruments). After gaining whole-cell access, cells were dialyzed for 5 minutes in voltage-clamp configuration at a command potential of −80mV. Membrane properties were assessed in current-clamp configuration as previously described [38] using a series of 1s current injections (−150pA to +500pA, Δ25pA) to evoke spike-firing.

For electrical- and optically-evoked IPSC recordings, the recording pipette was filled with a potassium-based internal solution (in mM: 125 K-gluconate, 4 NaCl, 10 HEPES, 0.1 EGTA, 4 MgATP, 0.3 Na_2_GTP, 10 Tris-phosphocreatine; pH adjusted to 7.3 with KOH, 296 mOsm). IPSCs were evoked using a paired-pulse protocol under voltage-clamp configuration at −60 mV. Excitatory currents were abolished using the AMPAR receptor blocker NBQX (10 μM) For electrically-evoked IPSCs, two pulses of electrical stimulation (0.14 ms, 100 ms ISI, 0.1 Hz) were delivered through a bipolar stimulating electrode placed in L1 of PL-PFC. In optogenetic experiments, GABA release was stimulated from ChR2-expressing neurons using two pulses of 470 nm blue light (1-2 ms, 100 ms ISI, 0.1 Hz) delivered via wide-field illumination through the microscope objective. Light intensity was adjusted to produce stable oIPSCs between 100-400 pA amplitude. Paired-pulse ratio (PPR) was calculated by dividing the average amplitude of the second IPSC over the average amplitude of the first during the 10-minute baseline and the last two minutes of drug washes. Coefficient of variance (CV) was calculated by dividing the standard deviation over the mean amplitude of the first IPSC during the last 5 minutes of the drug wash. sIPSCs were recorded under voltage-clamp configuration at −80 mV in NBQX with a cesium-based internal solution (in mM: 125 CsCl, 5 NaCl, 10 HEPES, 5 QX-314 bromide, 4 Mg-ATP, 0.2 Na-GTP, 10 Tris-phosphocreatine; pH adjusted to 7.3 with CsOH, 296 mOsm). Changes in access resistance (R_a_) were monitored throughout all recordings, and cells that showed an increase or decrease in R_a_ >20% were discarded and excluded.

### Viruses

| **Virus** | **Source** | **Stock Titer** |
| --- | --- | --- |
| AAV<sub>5</sub>-EF1a-DIO-hChR2(H134R)-EYFP WPRE | Addgene | $\geq 7e^{12}$ |
| AAV<sub>8</sub>-nEF-Coff/Fon-ChR2(ET/TC)-EYFP | Addgene | $\geq 1e^{13}$ |
| AAV<sub>5</sub>-mDlx-Flex-ChR2-mCherry | VectorBuilder | $1.63e^{13}$ |
| AAV<sub>1</sub>-Flex-tdTomato | Addgene | $\geq 1e^{13}$ |
| AAV<sub>8</sub>-EF1a-fDIO-mCherry | Addgene | $\geq 7e^{12}$ |
| AAV<sub>8</sub>-hDlx-Flex-GFP-Fishell-6 | Addgene | $\geq 7e^{12}$ |
| AAV<sub>5</sub>-mMORp-EYFP | Stanford | $1.1 e^{13}$ |

AAV_5_-EF1a-DIO-hChR2(H134R)-EYFP WPRE, AAV_8_-nEF-Coff/Fon-ChR2(ET/TC)-EYFP, AAV_5_-mDlx-Flex-ChR2-mCherry were used for Cre- or Flp-driven ChR2 expression in PV, SST, VIP, and CCK-INs. To record SST-IN transmission onto PV-INs, AAV_8_-nEF-Coff/Fon-ChR2(ET/TC)-EYFP and AAV_1_-Flex-tdTomato were combined 2:1. To record VIP-IN transmission onto fluorescently labeled SST-INs, AAV_8_-nEF-Coff/Fon-ChR2(ET/TC)-EYFP and AAV_8_-EF1a-fDIO-mCherry were combined 1:1. AAV_8_-hDlx-Flex-GFP was used to label CCK-INs. AAV_5_-mMORp-EYFP [65] were used to label MOR+ cells. AAV_5_-mMORp-EYFP was diluted 1:7 in cold phosphate-buffered saline.

### Stereotaxic Surgery

Stereotaxic surgery was performed in mice 5-6 weeks of age under aseptic conditions. Mice were initially anesthetized with 3% isoflurane and maintained on 1-2% for the duration of the surgery. Prior to the procedure, mice were administered 5 mg/kg carprofen dissolved in 0.9% saline. The scalp was cleaned with ethanol and betadine and an incision was made. A single burr hole was made above the PL-PFC (AP: +2.2, ML: +/- 0.3, DV: −1.8) and 0.4-0.6 μL of virus was delivered at 0.1 μL/minute by a glass capillary nanoinjector (Stoelting). The injector was left in place for 2 minutes at the termination of the infusion and then slowly retracted. The incision was closed with surgical glue (Vetbond) and treated with topical antibiotic ointment. Animals recovered on a heating pad in a clean cage before being returned to the homecage. Animals received post-operative injections of Carprofen (5 mg/kg) for 72 hours following the surgical procedure. At least 3 weeks was allowed for viral incubation before animals were sacrificed for electrophysiological recordings.

### Drugs

DAMGO ([D-Ala^2^,N-Me-Phe^4^,Glyol^5^]Enkephalin), forskolin, tetrodotoxin (TTX), and NBQX were purchased from HelloBio. DPDPE ([D-Pen2,5]Enkephalin) was purchased from Tocris. DAMGO, DPDPE, and TTX were dissolved in deionized water. Forskolin and NBQX were dissolved in DMSO. All drugs were made at 1000x, aliquoted, and frozen until use. DAMGO and DPDPE each have low-nanomolar binding affinities for MOR and DOR, respectively, in cell-based assays with >30-fold selectivity over other opioid receptors [81].

### Imaging

Viral targeting was confirmed with live imaging during slice recordings. Images were acquired with a 3-channel LED system (pE300) and a SciCam Pro CCD camera. Animals with mistargeted viral injections were discarded and excluded from analysis.

### Statistics

ClampFit 11.2 (Molecular Devices) was used for primary analysis of electrophysiological data. Statistical analyses were performed using GraphPad Prism (v10). The total number of animals and cells included in an experiment are denoted by “N” and “n”, respectively. For all analyses, p<0.05 was considered significant. Data were analyzed using two-tailed student’s t-test, one-sample t-test, two-way ANOVA, three-way ANOVA, repeated measures ANOVA, or mixed-effect analysis. Bonferroni post-hoc comparisons were used for analyses requiring multiple comparisons. Statistical analyses are reported in the legends and text. Data are presented as mean±SEM.

## Results

### MOR and DOR suppress spontaneous PL-PFC inhibitory transmission through different molecular mechanisms

Given their expression within inhibitory cells in PFC [12, 18, 36, 45, 73, 80], we first compared and contrasted how MOR and DOR modulate GABA transmission onto pyramidal cells. We made whole-cell recordings from pyramidal neurons (PNs) in L2/3 of the PL-PFC and recorded spontaneous inhibitory postsynaptic currents (sIPSCs). To activate MOR or DOR in separate cells, we applied either DAMGO ([D-Ala^2^, N-MePhe^4^, Gly-ol]-enkephalin) (MOR EC_50_=1.5 nM) [52] or DPDPE (d-Pen^2^,d-Pen^5^ enkephalin) (DOR EC_50_=5.2 nM) [17]. In each cell, we examined the sensitivity of sIPSC frequency and amplitude to 100 nM or 1 µM concentrations of each drug **(Figure 1A)**. Previous studies found that effects of 1 μM DAMGO and DPDPE in brain slices are blocked by MOR- and DOR-selective antagonists, respectively [2, 45]. We found that DAMGO and DPDPE suppressed sIPSC frequency **(Figure 1B-C)**, while sIPSC amplitude remained intact **(Figure 1D-E)**, consistent with a presynaptic locus of inhibition. Interestingly, equipotent concentrations of DPDPE were more effective at suppressing sIPSC frequency than DAMGO (DAMGO: 84 ± 5% of baseline vs DPDPE: 59 ± 4% of baseline) **(Figure 1C)**. One possible explanation for this difference is that DAMGO activates MORs on synapses that disinhibit IN activity. To test this idea, we repeated these experiments in the presence of tetrodotoxin (TTX) to block action potentials and isolate monosynaptic transmission. TTX did not alter the effects of DAMGO or DPDPE: the reductions in miniature IPSC (mIPSC) frequency were comparable to effects on sIPSCs from Figure 1C **(Figure 1F)**. mIPSC amplitude was also not affected (data not shown). While this observation does not rule out MOR/DOR actions at disinhibitory synapses, we hypothesized that a difference in molecular mechanism drives the differential suppression of sIPSC frequency by MOR and DOR.

**Figure 1.**
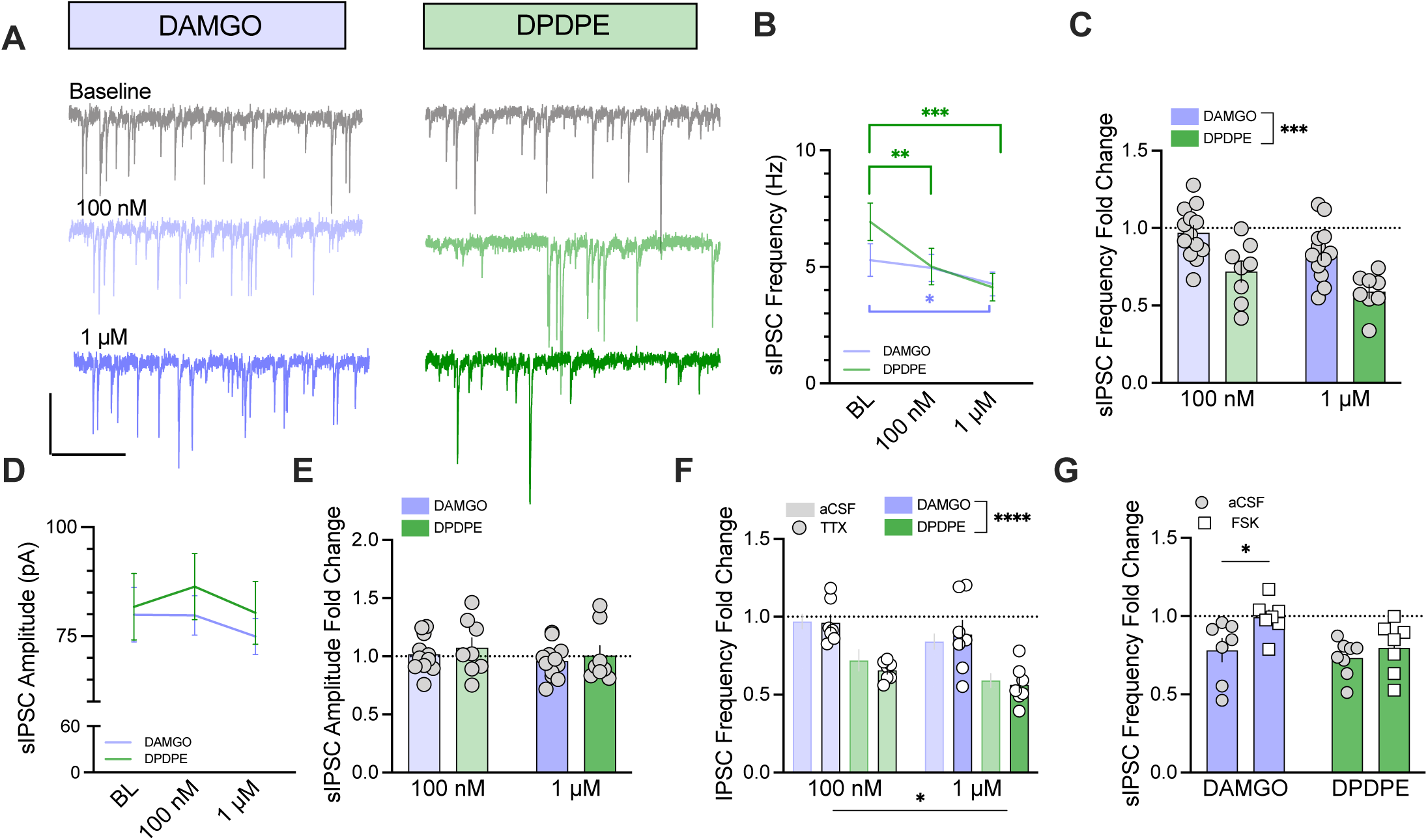
MOR and DOR suppress PL-PFC spontaneous inhibitory transmission. **A)** Representative traces of spontaneous inhibitory postsynaptic currents (sIPSCs) in pyramidal cells at baseline (grey) and during bath-application of 100 nM and 1 μM DAMGO (light purple and dark purple) or DPDPE (light green and dark green). Scale bar represents 100 pA and 1 second. **B)** DAMGO and DPDPE decrease sIPSC frequency (two-way RM ANOVA, agonist x concentration interaction: F_2.38_=8.6, p= 0.0008). DAMGO: BL= 5.3 ± 0.7 Hz, 100 nM = 5.0 Hz ± 0.6 Hz, 1 μM= 4.3 ± 0.5 Hz; DPDPE: BL= 6.9 ± 0.8 Hz, 100 nM= 5.0 ± 0.8 Hz, 1 μM = 4.1 ± 0.6 Hz. sIPSCs recorded from n/N= 8-13/6-8 cells/mice. **C)** DPDPE suppresses sIPSC frequency to a greater extent than DAMGO (two-way RM ANOVA, main effect of agonist: F_1,19_= 16.1, p = 0.0007, main effect of concentration: F_1,19_= 9.4, p = 0.0063). DAMGO: 100 nM= 1.0 ± 0.05, 1 μM= 0.8 ± 0.05; DPDPE 100 nM= 0.7 ± 0.07, 1 μM: 0.6 ± 0.04. **D)** DAMGO and DPDPE do not affect sIPSC amplitude (two-way RM ANOVA, main effect of agonist: p= 0.5615, main effect of concentration: p= 0.25). DAMGO: BL= 80 ± 6.2 pA, 100 nM= 80 ± 4.5 pA, 1 μM= 75 ± 4.1 pA; DPDPE: BL= 82 ± 7.6 pA, 100 nM= 86 ± 7.6 pA, 1μM= 80 ± 7.2 pA. **E)** There is no difference between the effect of DAMGO and DPDPE on sIPSC amplitude (two-way RM ANOVA, main effect of agonist: p= 0.51, main effect of concentration: p= 0.11). DAMGO: 100 nM= 1.0 ± 0.04, 1 μM = 1.0 ± 0.04; DPDPE: 100 nM= 1.1 ± 0.08, 1μM= 1.0 ± 0.08. sIPSCs recorded from n/N= 8-13/6-8 cells/mice. **F)** DAMGO and DPDPE have similar effects on sIPSCs and miniature IPSCs (mIPSCs) (three-way RM ANOVA, main effect of agonist: F_1,31_= 34.6, p<0.0001, main effect of concentration: F_1,31_= 12.1, p= 0.0015, main effect of TTX: p= 0.77). DAMGO mIPSC: 100 nM= 1.0 ± 0.05, 1 μM= 0.9 ± 0.09; DPDPE mIPSC: 100 nM= 0.7 ± 0.02, 1 μM= 0.6 ± 0.05. mIPSCs recorded from n/N= 7/3-4 cells/mice, control data from **1C**. **G)** Forskolin (FSK) blocks the effect of DAMGO but not DPDPE (two-way ANOVA, main effect of forskolin: F_1,25_= 6.0 p= 0.0218, main effect of agonist: F_1,25_= 4.7 p= 0.0391). Control DAMGO= 0.8 ± 0.07, FSK+DAMGO= 1.0 ± 0.04; Control DPDPE= 0.7 ± 0.04, FSK+DPDPE= 0.8 ± 0.06. n/N= 7-8/4-6 cells/mice. *p<0.5, **p<0.01, ***p<0.001, Bonferroni’s multiple comparisons test.

Opioid receptors engage canonical Gα_i/o_-dependent signaling to initiate chemical messaging. Opioid receptor activation attenuates adenylyl cyclase (AC) function to reduce cAMP levels, impacting downstream effectors and ion channels [64]. We probed this pathway by using the AC activator forskolin (FSK) to override Gα_i/o_ signaling. Concurrent application of FSK (10 µM) blocked the DAMGO-induced suppression of sIPSC frequency but did not prevent inhibition by DPDPE (DAMGO control: 78 ± 7% of baseline vs FSK: 99% ± 4% of baseline; Control DPDPE: 73 ± 4% of baseline vs FSK: 80 ± 6% of baseline) **(Figure 1G)**. These results show that MOR suppresses spontaneous inhibitory transmission via a cAMP-dependent mechanism and provide molecular insight into the differential efficacy of MOR and DOR in reducing GABA release probability.

### MOR and DOR suppress evoked PL-PFC inhibitory transmission through dissociable mechanisms

Synaptic transmission involves both spontaneous and action potential-dependent release. The processes mediating spontaneous and evoked neurotransmission are partially segregated at inhibitory synapses [34], and this separation may occur through multiple molecular mechanisms. MOR signaling reduces evoked IPSCs in orbitofrontal cortex (OFC) [45] and anterior cingulate cortex (ACC) [12], and DOR reduces evoked IPSCs in L5 of the mPFC [2], but results from comparative studies have not been reported. Here, we directly compared MOR and DOR suppression of evoked IPSCs in L2/3 PL-PFC. We delivered paired-pulse electrical stimulation to L1 of PL-PFC and recorded electrically-evoked IPSCs (eIPSCs) in L2/3 PNs **(Figure 2A)**. DAMGO (1 µM, 5 min) caused an acute suppression of eIPSC amplitude that gradually reversed upon washout **(Figure 2B-C)**. We also monitored paired-pulse ratio (PPR) as a proxy for changes in presynaptic neurotransmitter release probability. We observed a modest increase in PPR that emerged following the drug washout (min 40-45), consistent with a reduction in GABA release probability **(Figure 2D)**. DPDPE (1 µM, 5 min) also suppressed eIPSC amplitude **(Figure 2E-F)** and increased PPR at the long-term timepoint **(Figure 2G)**. Like with spontaneous transmission, DAMGO and DPDPE displayed different efficacies for suppressing eIPSCs; however, by contrast, MOR activation suppressed eIPSC amplitude to a greater extent than DOR (DAMGO: 68 ± 3% of baseline vs DPDPE: 79 ± 4% of baseline) **(Figure 2H)**.

**Figure 2.**
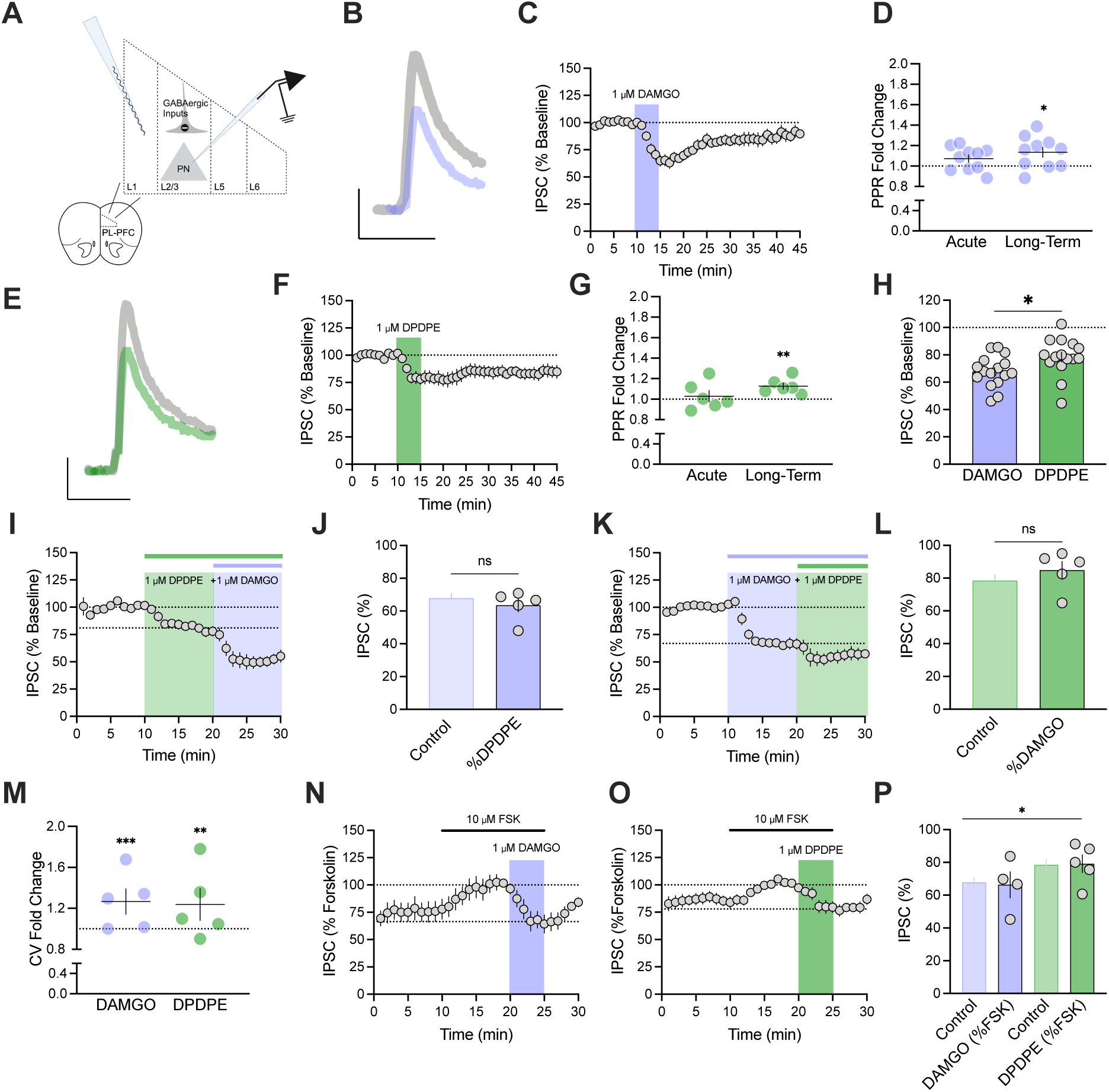
MOR and DOR inhibition of evoked transmission is additive. **A)** Schematic depicting experimental preparation for evoked IPSC (eIPSC) recordings. **B)** Representative traces of eIPSC amplitude at baseline (grey trace) and during the last 2 minutes of 1 μM DAMGO application (purple trace). Scale bar represents 50 pA and 50 ms. **C)** DAMGO acutely suppresses eIPSC amplitude. Timecourse plot shows eIPSC amplitude expressed as a percentage of baseline eIPSC amplitude. **D)** Change in paired-pulse ratio (PPR) is significant at long-term but not acute timepoints (one-sample t-tests, acute: p= 0.0776, long-term: p= 0.0232). Acute= 1.1 ± 0.03, long-term= 1.1 ± 0.05. n/N= 10/7 cells/mice. **E)** Representative traces of eIPSC amplitude at baseline (grey trace) and during the last 2 minutes of 1 μM DPDPE application (green trace). Scale bar represents 50 pA and 50 ms. **F)** DPDPE suppresses eIPSC amplitude. Timecourse plot shows eIPSC amplitude expressed as a percentage of baseline eIPSC amplitude. **G)** Change in PPR is significant at long-term but not acute timepoints (one-sample t-tests, acute: p= 0.6128, long-term: p= 0.0087). Acute= 1.0 ± 0.05, long-term= 1.1 ± 0.03. n/N= 6/3 cells/mice. **H)** DAMGO causes a greater acute suppression of eIPSC amplitude than DPDPE (unpaired t-test, p= 0.0260). DAMGO: 68 ± 2.8 %; DPDPE: 79 ± 3.5 %. n/N= 6-10/3-7 cells/mice. **I)** Timecourse plot of eIPSC amplitude expressed as a percentage of baseline eIPSC amplitude during sequential application of DPDPE and DAMGO. **J)** Pretreatment with DPDPE does not prevent eIPSC suppression by DAMGO (unpaired t-test p= 0.4492). The effect of DAMGO is expressed as a percentage of average eIPSC amplitude during the DPDPE wash. % DPDPE: 64 ± 4.1 %. n/N= 5/4 cells/mice, control data from **2H**. **K)** Timecourse plot showing eIPSC amplitude expressed as a percentage of baseline eIPSC amplitude during sequential application of DAMGO and DPDPE. **L)** Pretreatment with DAMGO does not prevent eIPSC suppression by DPDPE (unpaired t-test, p= 0.3716). The effect of DPDPE is expressed as a percentage of average eIPSC amplitude during the DAMGO wash. % DAMGO: 85 ± 5.4 %. n= 5/5 cells/mice, control data from **2H**. **M)** Extended DAMGO and DPDPE treatment increased coefficient of variance (CV) (one-sample t-tests, DAMGO: p= 0.0005, DPDPE: p= 0.0013). DAMGO= 1.3 ± 0.1; DPDPE= 1.2 ± 0.2. n= 5/4-5 cells/mice per group. **N)** Timecourse plot of eIPSC amplitude expressed as a percentage of baseline eIPSC amplitude during FSK and DAMGO application. **O)** Timecourse plot of eIPSC amplitude expressed as a percentage of baseline eIPSC amplitude during FSK and DPDPE application. **P)** FSK does not prevent eIPSC suppression by DAMGO or DPDPE (two-way ANOVA, main effect of agonist: F_1,36_= 6.4, p= 0.0156, main effect of forskolin: p= 0.9824). The effect of DAMGO or DPDPE with FSK is expressed as a percentage of average eIPSC amplitude during FSK pretreatment. Control DAMGO= 68 ± 2.8 %, FSK+DAMGO= 66 ± 7.7 %; Control DPDPE= 78 ± 3.5 %, FSK+DPDPE= 80 ± 5.0 %. n/N= 4-5/4-5 cells/mice, control data from **2H**. *p<0.05

To test whether MOR and DOR suppress evoked GABA release through a shared mechanism, we applied DAMGO and DPDPE sequentially in an occlusion experiment. If MOR and DOR use shared machinery in common cell types, then activation of one receptor system should preclude further suppression of eIPSC amplitude by the other. In these experiments, we extended drug application times to 10 minutes to ensure stable suppression of eIPSCs. In the presence of DPDPE, the addition of DAMGO further decreased eIPSC amplitude to a similar magnitude as seen under previous control conditions (64 ± 4% of baseline vs 68 ± 3% of baseline) **(Figure 2I-J)**.

Conversely, pre-application of DAMGO did not reduce subsequent actions of DPDPE (85 ± 4% of baseline vs 79 ± 4% of baseline) **(Figure 2K-L)**. In both experiments, DAMGO and DPDPE (1 µM, 10 min) alone increased the coefficient of variance (CV), providing additional corroboration of a change in presynaptic release probability **(Figure 2M)**. These data indicate that MOR and DOR suppress PFC inhibitory transmission through separate mechanisms in L2/3 of PL-PFC.

Given that forskolin blocked the suppression of spontaneous transmission by MOR but not DOR **(Figure 1G)**, we hypothesized that the greater effects of DAMGO on eIPSC amplitude were driven by cAMP signaling. The results did not support this hypothesis, as pre-treatment with FSK (10 µM) had no effect on either DAMGO- or DPDPE-induced suppression of eIPSC amplitude **(Figure 2N-P)**. Taken together, these data suggest that MOR and DOR reduce spontaneous and evoked transmission through separate molecular mechanisms. Furthermore, the additive effects of DAMGO and DPDPE suggest that MOR and DOR may attenuate GABAergic transmission through distinct populations of INs.

### MOR and DOR differentially inhibit cortical IN output

Opioid receptor signaling can have diverse outcomes depending on which cells harbor that receptor. We used cell type-specific *ex vivo* optogenetics to test the hypothesis that MOR and DOR differentially modulate GABA release across IN subtypes.

Previous studies found that DOR activation potently suppresses inhibitory transmission from parvalbumin-expressing INs (PV-INs) onto PFC PNs [3, 12], while MOR-dependent modulation of PV-IN transmission is more complex, showing differential sensitivity to DAMGO depending on cortical subregion [12, 45]. To compare the effects of MOR and DOR signaling on PV-IN-mediated transmission, we examined the response of optically-evoked IPSCs (oIPSCs) from PV-INs onto pyramidal cells to DAMGO or DPDPE **(Figure 3A)**. MOR activation by DAMGO had no effect on oIPSC amplitude (95 ± 2% of baseline), while DPDPE potently suppressed PV-IN oIPSCs (60 ± 4% of baseline) **(Figure 3B-C)**. These results suggest that opioid-mediated suppression of PV-IN inhibitory transmission is primarily mediated through DOR expression at the presynaptic terminal.

**Figure 3.**
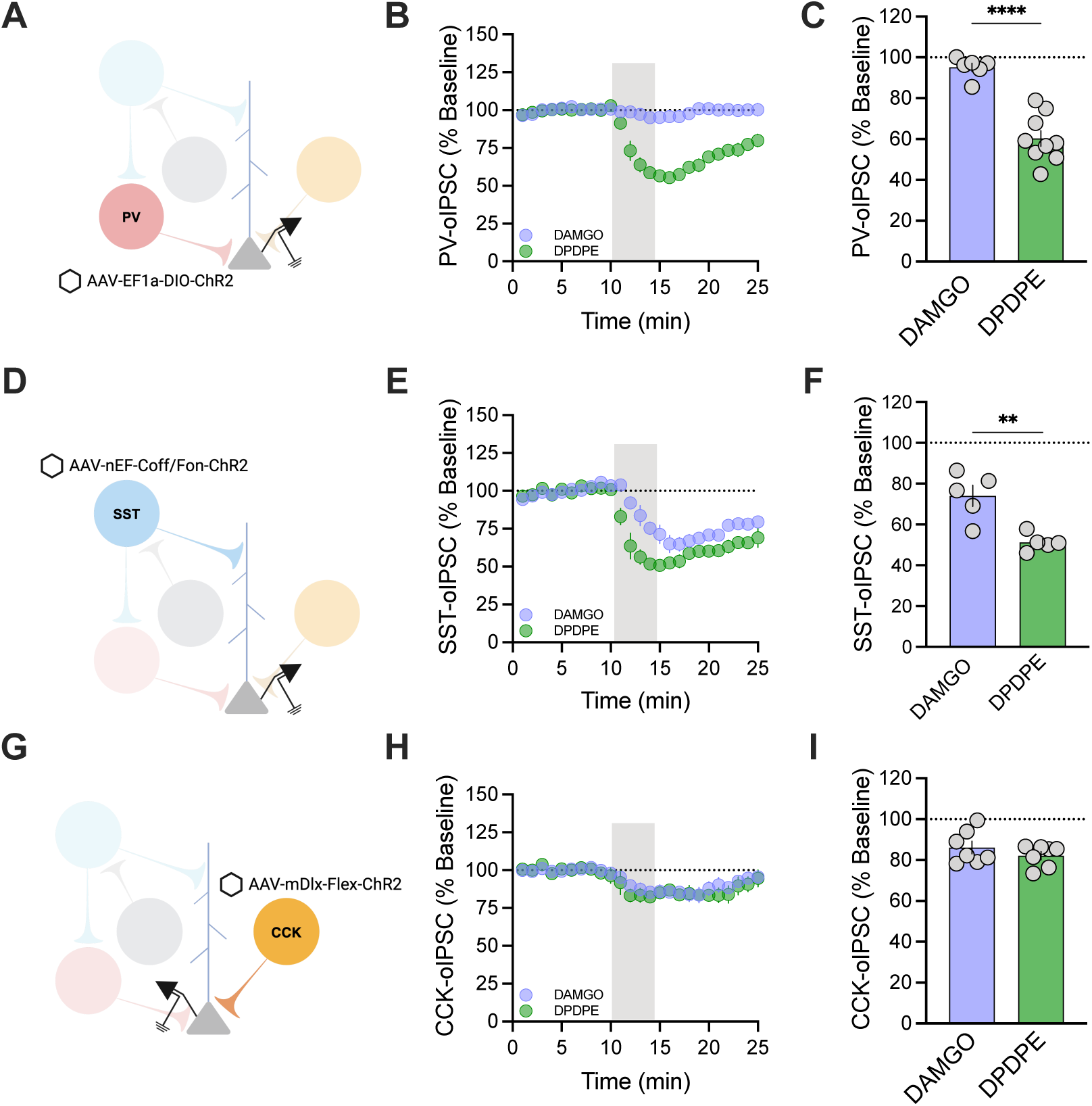
MOR and DOR differentially regulate cortical IN output. **A)** Schematic of viral construct and recording preparation used to record PV-IN optically-evoked IPSCs (oIPSCs). **B)** DPDPE suppresses PV-oIPSCs. Timecourse plot shows oIPSC amplitude during DAMGO or DPDPE application expressed as a percentage of baseline oIPSC amplitude. **C)** DPDPE causes a greater suppression of PV-IN output than DAMGO. (unpaired t test, p<0.0001). DAMGO: 95 ± 2.1 %; DPDPE: 60 ± 3.9 %. n/N= 6-9/8 cells/mice per group. **D)** Schematic of viral construct and recording preparation used to record SST-oIPSCs. **E)** DAMGO and DPDPE suppress SST-oIPSCs. Timecourse plot showing oIPSC amplitude during DAMGO or DPDPE application expressed as a percentage of baseline oIPSC amplitude. **F)** DPDPE causes a greater suppression of SST-IN output than DAMGO. (unpaired t test, p= 0.0033). DAMGO: 74 ± 5.2 %; DPDPE: 51 ± 1.9 %. n/N= 5/8 cells/mice per group. **G)** Schematic of viral construct and recording preparation used to record CCK-oIPSCs. **H)** DAMGO and DPDPE suppress CCK-oIPSCs. Timecourse plot showing oIPSC amplitude during DAMGO or DPDPE application expressed as a percentage of baseline oIPSC amplitude. **I)** DAMGO and DPDPE suppress CCK-IN output to similar levels (unpaired t test, p= 0.2939). DAMGO: 86 ± 3.1 %; DPDPE: 82 ± 2.0 %. n/N= 7/8 cells/mice per group.

Based on our previous observations that DAMGO potently suppresses eIPSCs in PL cortex, the insensitivity of PV-oIPSCs to DAMGO suggests that MOR likely suppress inhibitory transmission from other INs with higher efficacy. Somatostatin-expressing INs (SST-INs) in the rodent neocortex contain high levels of *Oprm1* and *Oprd1,* encoding for MOR and DOR, respectively [69]. Bath application of DPDPE inhibits SST-IN output onto pyramidal cells in L5 of the PL-PFC [2], but the extent to which MOR and DOR inhibit SST-IN transmission has not been compared. DAMGO and DPDPE both suppressed SST-IN oIPSC amplitude, though DPDPE showed higher efficacy than DAMGO (DAMGO: 74 ± 5% of baseline vs DPDPE: 51 ± 2% of baseline) **(Figure 3D-F)**.

INs that express cholecystokinin (CCK-INs) are also abundant in L2/3 PL-PFC and transmit monosynaptic IPSCS onto pyramidal cells [41]. The extent to which CCK-INs express functional opioid receptors has not been reported. Here, we found that DAMGO and DPDPE each modestly suppressed CCK-IN output (DAMGO: 86 ± 3% of baseline vs DPDPE: 82 ± 2% of baseline) **(Figure 3G-I)**, demonstrating that MOR and DOR expression on CCK-IN presynaptic terminals can regulate CCK-IN synaptic transmission. Ultimately, DOR is positioned to primarily control GABA release from L2/3 PL-PFC PV and SST-INs onto PNs, while MOR and DOR comparably suppress CCK-IN output.

### MOR and DOR regulate PFC disinhibitory circuits

Disinhibition plays a key role in behavior and memory processing by dampening the activity of other INs and enhancing pyramidal cell activity [6, 20, 79]. To our knowledge, there are no published studies examining opioid actions at disinhibitory synapses in PFC. Given that MOR and DOR potently suppress oIPSCs from SST-INs onto PNs **(Figure 3E-F)**, we next explored how MOR and DOR regulate SST-IN disinhibitory transmission. SST-INs can disinhibit PFC activity through connections with PV-INs [20]. To isolate inhibitory transmission from SST-INs onto PV-INs, we delivered a viral cocktail to express Cre-Off/Flp-On ChR2 vector and Cre-dependent tdTomato in the PL-PFC of double transgenic SST-Flp/PV-Cre mice **(Figure 4A)**. Both DAMGO and DPDPE suppressed SST-IN output onto PV-INs to approximately 70% of baseline **(Figure 4B)**. Interestingly, the effect of DPDPE was more modest at SST-PV synapses compared to SST-PN (71 ± 6% of baseline vs 51 ± 2% of baseline) **(Figure 4C)**, despite no significant difference in basal release probability across synapses **(Figure 4D)**.

**Figure 4.**
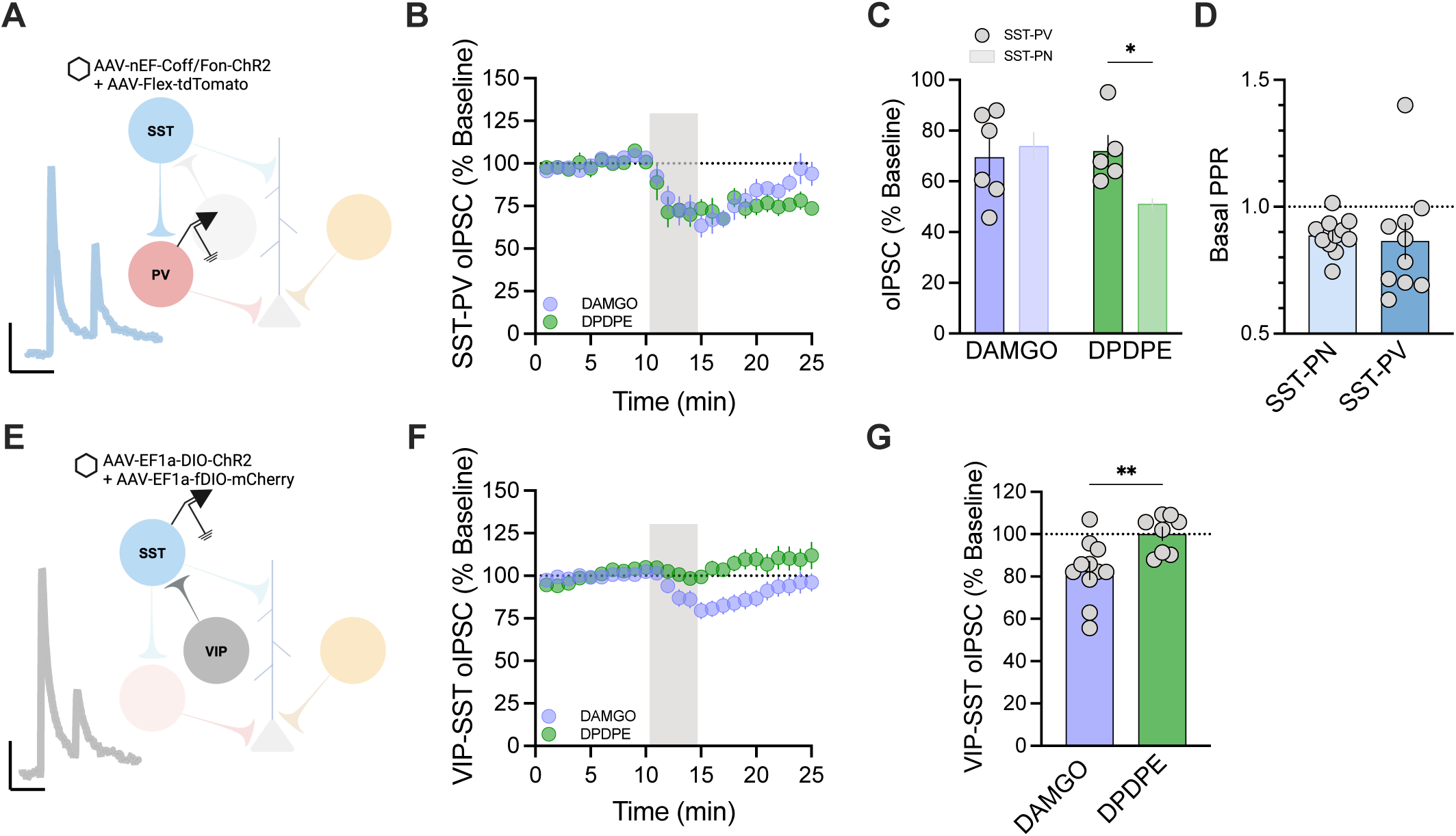
MOR and DOR regulate disinhibitory circuitry. **A)** Schematic of viral constructs and recording preparation used to record SST-oIPSCs onto PV-INs and representative traces. Scale bar represents 20 pA and 100 ms. **B)** SST-PV-oIPSCs are suppressed by DAMGO and DPDPE. Timecourse plot showing oIPSC amplitude during DAMGO or DPDPE application expressed as a percentage of baseline oIPSC amplitude. **C)** DAMGO and DPDPE suppress inhibitory transmission from SST-INs onto PV-INs. DPDPE is more effective at suppressing SST-oIPSCs onto pyramidal cells than PV-INs (two-way ANOVA, agonist x synapse interaction: F_1,17_= 4.9, p= 0.0409). SST-PV: DAMGO: 70 ± 7.1 %; DPDPE: 72 ± 6.1 %; SST-PN: DAMGO: 74 ± 5.2 %; DPDPE: 51 ± 1.9 %.SST-PN data from **3F**. **D)** SST-INs show similar basal release probability across SST-PV and SST-PN synapses (unpaired t-test, p= 0.7650). SST-PV PPR: 0.87 ± 0.071; SST-PN PPR: 0.89 ± 0.021. n= 5-6 cells/group from 5 mice. **E)** Schematic of viral constructs and recording preparation used to record VIP-oIPSCs in SST-INs and representative traces. Scale bar represents 20 pA and 100 ms. **F)** Timecourse plot showing oIPSC amplitude during DAMGO or DPDPE application expressed as a percentage of baseline oIPSC amplitude. **G)** DAMGO causes a greater suppression of VIP-IN output onto SST-INs than DAMGO. (unpaired t-test, p= 0.0074). DAMGO: 83 ± 4.3 %; DPDPE: 100 ± 3.1 %. n/N= 8-11/6 cells/mice per group.

INs that express vasoactive intestinal peptide (VIP-INs) form prominent disinhibitory circuits with SST-INs [44, 47, 61]. At VIP-SST synapses, we found that DAMGO modestly inhibited VIP-oIPSCs while DPDPE had no effect (DAMGO: 83 ± 4% of baseline vs DPDPE: 100 ± 3% of baseline) **(Figure F-G)**. These results show that MOR and DOR regulate primary synapses onto glutamatergic cells, as well as disinhibitory synapses between INs, to modulate cortical function.

### Opioids differentially regulate IN subtypes depending on receptor type and localization

In addition to presynaptic opioid receptors modulating neurotransmitter release probability, somato-dendritic opioid receptors can reduce cell excitability by activating G protein-coupled inwardly rectifying K^+^ (GIRK) channels [46, 70, 72, 77]. To assess this function, we recorded directly from PV, SST, CCK, and VIP-INs and monitored changes in holding current in response to DAMGO or DPDPE application. We patched fluorescently identified PV, SST, and VIP-INs from genetically engineered PV-Cre-tdTomato, SST-Flp-tdTomato, or VIP-Cre-tdTomato reporter mice, respectively. PV-INs and SST-INs responded to DAMGO and DPDPE, but both cell types showed a larger change in holding current with DPDPE (PV-IN: 21 ± 3 pA vs 7 ± 2 pA; SST-IN: 27 ± 11 pA vs 12 ± 3 pA) **(Figure 5A)**. VIP-IN holding current did not change with either drug **(Figure 5A)**.

**Figure 5.**
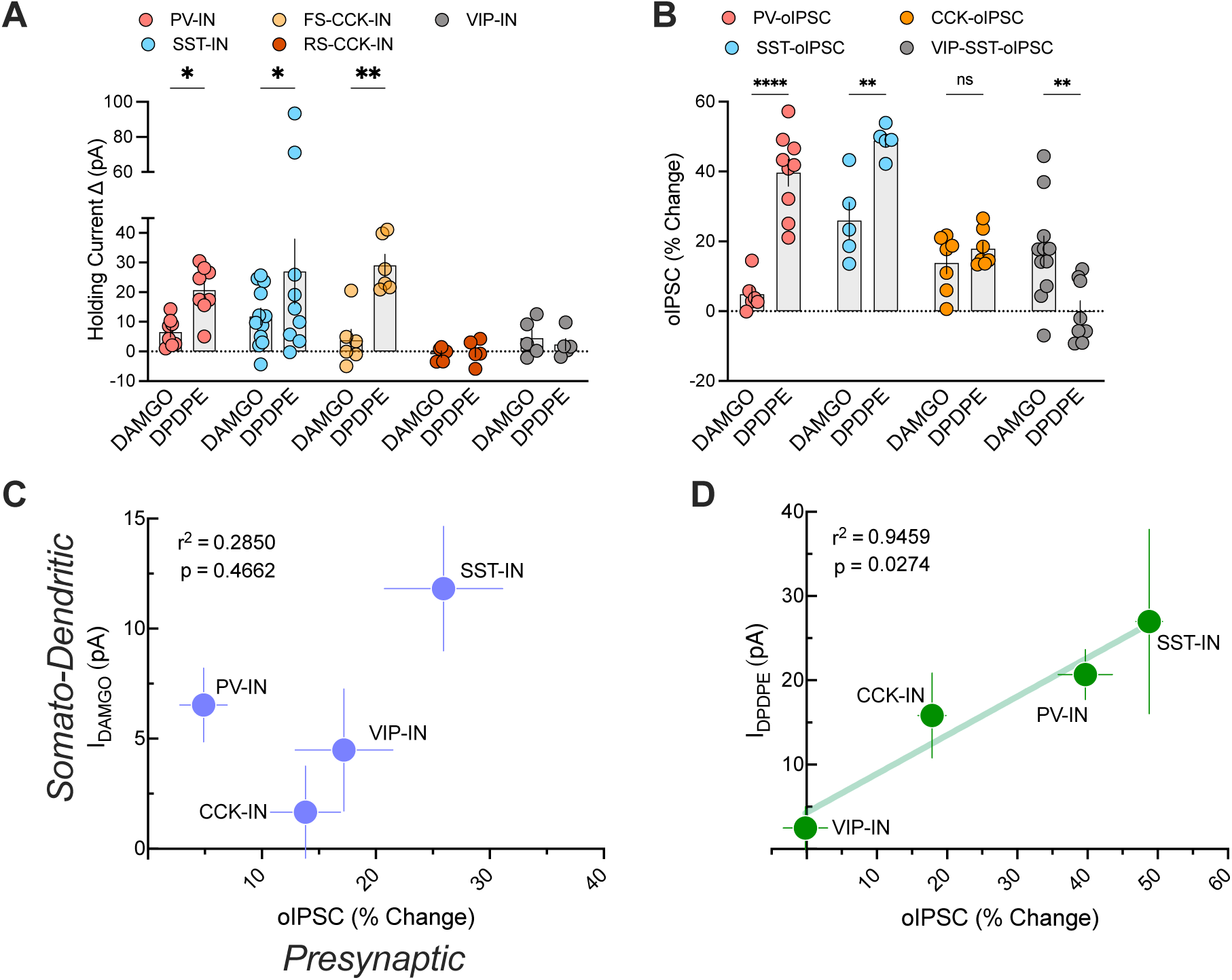
Pre- and postsynaptic opioid regulation of inhibitory transmission varies by cell and receptor type. **A)** Summary of the maximal DAMGO or DPDPE-induced change in holding current. MOR- and DOR-induced change in holding current varies across IN types (two-way ANOVA, main effect of cell type: F_4,59_= 4.8, p= 0.0022, main effect of agonist: F_1,59_= 9.3, p= 0.0035; uncorrected Fisher’s LSD *p<0.05, **p<0.01). PV-IN: DAMGO: 6.5 ± 1.7 pA; DPDPE: 21 ± 3 pA; SST-IN: DAMGO: 12 ± 2.8 pA; DPDPE: 27 ± 11 pA %; FS-CCK-IN: DAMGO: 3.8 ± 3.6 pA; DPDPE: 29 ± 3.7 pA; RS-CCK-IN: DAMGO: −0.9 ± 1.1 pA; DPDPE: −0.04 ± 1.8 pA; VIP-IN: DAMGO: 4.5 ± 2.8 pA; DPDPE: 2.3 ± 2 pA. n/N = 5-12/3-6 cells/mice per group. **B)** Summary of the maximal DAMGO or DPDPE-induced change in oIPSC. MOR- and DOR-induced suppression of oIPSC varies across IN types (two-way ANOVA, agonist x synapse interaction: F_4,59_= 13.11, ****p<0.0001; Bonferroni’s multiple comparisons test **p<0.01, ****p<0.0001). PV-IN: DAMGO: 4.9 ± 2.1 %; DPDPE: 40 ± 3.9 %; SST-IN: DAMGO: 26 ± 5.2 %; DPDPE: 49 ± 1.9 %; CCK-IN: DAMGO: 14 ± 3.1 %; DPDPE: 18 ± 2.1 %; VIP-IN: DAMGO: 17 ± 4.3 %; DPDPE: −0.2 ± 3.1 %. n/N = 5-11/5-6 cells/mice per group. **C)** The relationship between DAMGO-induced change in holding current and change in oIPSC is nonlinear (simple linear regression, R^2^= 0.2850, p= 0.4662). Vertical error bars represent holding current change SEM and horizontal error bars represent oIPSC change SEM. **D)** The relationship between DPEPE-induced change in holding current and change in oIPSC is linear (simple linear regression, R^2^= 0.9459, p= 0.0274). Vertical error bars represent holding current change SEM and horizontal error bars represent oIPSC change SEM.

To selectively label CCK-INs, we delivered Cre-dependent GFP under the Dlx enhancer element into the PL-PFC of CCK-Cre-VGAT-Flp mice [22, 41]. Using this approach, we observed two distinct populations of CCK-INs previously characterized by Kamalova et al. [41]. Approximately half of the total GFP+ CCK cells that we patched (15/34) displayed a fast-spiking phenotype (FS-CCK-INs), which we defined as having a firing frequency ≥70 Hz. The remaining cells were classified as regular-spiking CCK-INs (RS-CCK-INs). We applied DAMGO and DPDPE (1 μM, 5 min) to a subset of FS- and RS-CCK-INs while holding cells at a command potential of −60 mV to enhance the driving force for potassium. FS-CCK-INs displayed large hyperpolarizing outward currents in response to DPDPE (29 ± 4 pA), consistent with enhanced GIRK channel conductance, and no significant response to DAMGO **(Figure 5A)**. To account for GIRK channel desensitization [63], we quantified the peak response during the 5 minute drug wash. There was no effect of DAMGO or DPDPE on RS-CCK-INs, suggesting that DOR cellular expression patterns may be cell subtype-specific.

For DPDPE, the magnitude of the somato-dendritic response generally coordinated with the presynaptic response. Cells that showed outward currents upon DOR activation **(Figure 5A)** also showed a reduction in their synaptic output **(Figure 5B)**. We observed a more complex relationship between pre- and postsynaptic MOR function. The relationship between the change in holding current and reduction in oIPSC amplitude was nonlinear for DAMGO **(Figure 5C)**, suggesting there are cell type-specific differences in the regulation of presynaptic and somato-dendritic MOR. By contrast, we observed a strong correlation (r^2^ = 0.9459) between the change in holding current and reduction in oIPSC amplitude for DOR **(Figure 5D)**. Together, these observations suggest there are profound differences in subcellular functions of MOR but not DOR in PFC INs.

### MOR expression is predominantly localized to fast-spiking interneurons

Our finding that pre- and post-synaptic functions of MOR are not correlated within PFC INs raised concerns that (1) our selected measurements of MOR function are limited and incomplete, and (2) these measurements may not provide clear insight into overall, cellular expression patterns. To address these two limitations, we used a novel, orthogonal approach to characterize MOR expression in PFC neurons based on expression of *Oprm1*. To label MOR+ cells, we delivered AAV_5_-mMORp-EYFP into the PL-PFC of congenic C57BL/6J mice [65]. Three weeks following the viral infusion, we performed current-clamp experiments from EYFP+ cells in L2/3. From these experiments, we collected a series of membrane properties including a spike-firing input-output curve and rheobase, which we used in conjunction with cell capacitance (C_m_) to classify cells as either putative excitatory PNs or INs. We were able to further subdivide putative GABAergic cells into groups that had membrane properties characteristic of low-threshold spiking INs (LTS), fast-spiking INs (FSI), and regular-spiking non-pyramidal neurons (RSNP) **(Figure 6A).** These criteria were applied *post hoc* to fluorescently identified PV-INs, SST-INs, CCK-INs, and VIP-INs, confirming that these parameters accurately capture membrane physiology representative of FSIs, LTS cells, and RSNP cells **(Figure 6B-E).** Of the 33 MOR-EYFP+ cells that we patched, 15 had membrane properties characteristic of FSIs (45%), 11 appeared characteristic of RSNP neurons (33%), 6 appeared characteristic of LTS neurons (18%), and 1 cell was classified as a putative pyramidal cell (3%) **(Figure 6G)**. Cells identified by EYFP fluorescence displayed outward currents in response to DAMGO (1 μM) **(Figure 6H)**, confirming functional MOR expression. Using this promoter-based tool to complement our physiological assessment of MOR function in PFC INs, we show that FSIs and RSNP cells comprise the majority of MOR+ neurons in the PL-PFC, despite relatively modest effect sizes for functional assessments of MOR signaling.

**Figure 6.**
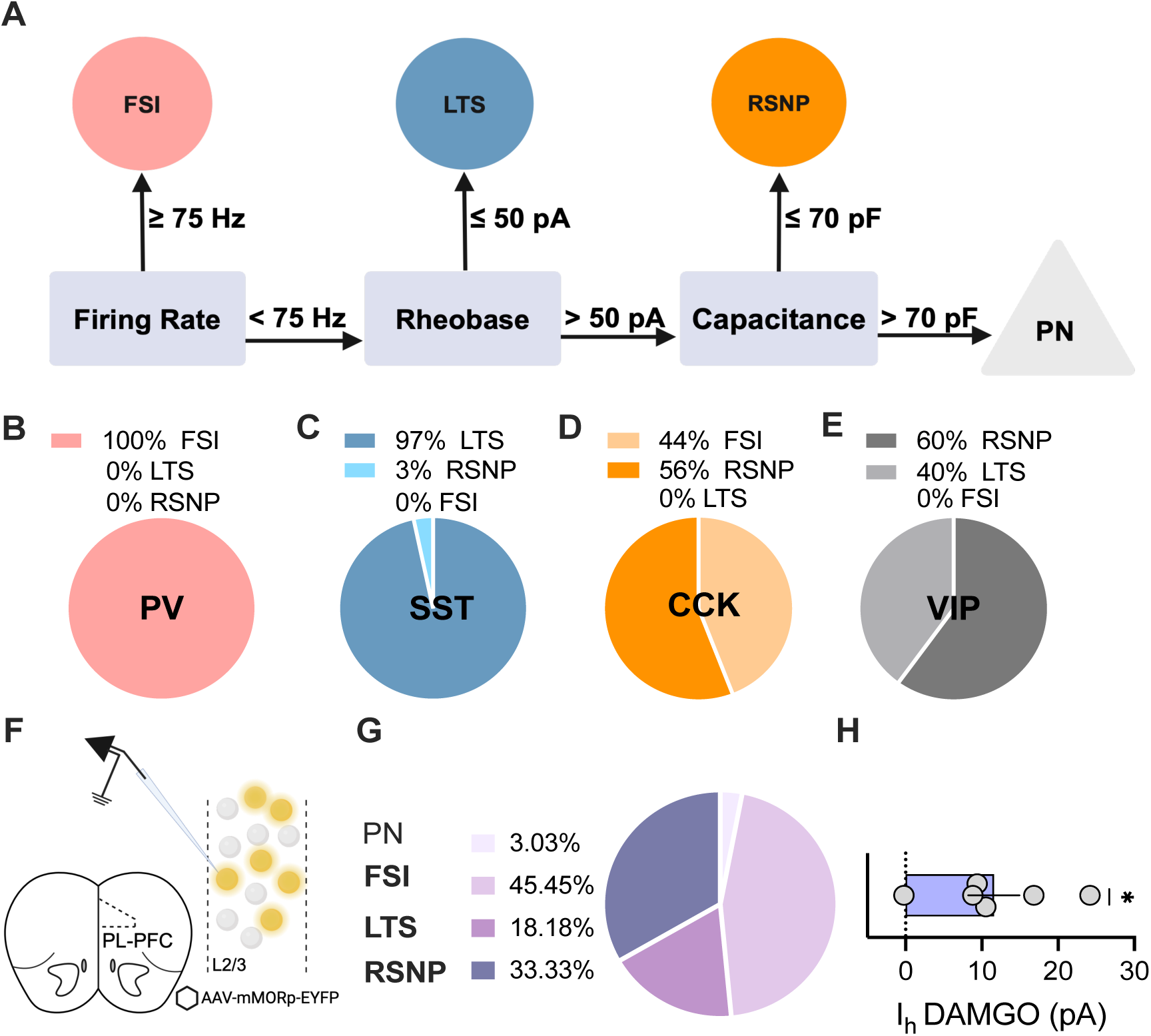
Cellular expression patterns of MOR in L2/3 of PL-PFC. **A)** Schematic depicting workflow used for cell type classification. **B)** Proportion of tdTomato+ PV-INs that displayed membrane physiology characteristic of low-threshold spiking (LTS) cells, regular-spiking non-pyramidal (RSNP) cells, and FSIs. n= 25 total cells. **C)** Proportion of tdTomato+ SST-INs that displayed membrane physiology characteristic of low-threshold spiking (LTS) cells, regular-spiking non-pyramidal (RSNP) cells, and FSIs. n= 30 total cells. **D)** Proportion of GFP+ CCK-INs that displayed membrane physiology characteristic of low-threshold spiking (LTS) cells, regular-spiking non-pyramidal (RSNP) cells, and FSIs. n= 34 total cells. **E)** Proportion of tdTomato+ VIP-INs that displayed membrane physiology characteristic of low-threshold spiking (LTS) cells, regular-spiking non-pyramidal (RSNP) cells, and FSIs. n= 10 total cells. **F)** Schematic of viral construct and recording preparation for *MORp*-EYFP+ cells. **G)** Proportion of EYFP+ cells that displayed membrane physiology characteristic of pyramidal neurons (PNs), FSIs, LTS neurons, and RSNP neurons. **H)** EYFP+ cells exhibit outward currents upon MOR activation (one-sample t-test, *p<0.05).

## Discussion

In this study, we show that MOR and DOR differentially regulate inhibitory transmission onto L2/3 PL-PFC pyramidal cells depending on mode of inhibition, inhibitory input, and putative receptor localization. MOR-mediated suppression of spontaneous inhibitory synaptic transmission is cAMP-dependent, while DOR suppresses sIPSCs independently of cAMP signaling. MOR and DOR regulate action potential-dependent transmission through dissociable mechanisms that suppress eIPSCs with different efficacy. We demonstrate that MOR and DOR regulate PFC GABA signaling through peri-somatic *and* presynaptic regulation of IN activity, though there is significant heterogeneity in opioid sensitivity across inhibitory synapses. Our findings emphasize the complexity of endogenous opioid signaling in PFC microcircuitry and highlight important considerations for interpreting opioid actions within intact circuitry.

### Differential suppression of spontaneous and evoked transmission by MOR and DOR

Consistent with the existing literature [14, 35, 49, 50, 55], we show that MOR and DOR suppress spontaneous transmission. DAMGO and DPDPE reduced sIPSC frequency but not amplitude, consistent with a presynaptic inhibition of GABA release. DPDPE inhibited sIPSC frequency to a greater extent than DAMGO, indicating that MOR and DOR regulate spontaneous transmission through different mechanisms. Indeed, enhancing cAMP signaling with FSK blocked MOR-mediated suppression of sIPSC frequency but not did not alter the effect of DPDPE. Unlike spontaneous transmission, MOR- and DOR-induced suppression of stimulation-evoked inhibitory transmission was not sensitive to FSK treatment. This observation contrasts with previous reports of cAMP-dependent MOR- and DOR-induced long-term depression of inhibitory transmission in other cortical [45] and subcortical [59] structures. One interpretation for these data is that MOR and DOR can situationally engage signaling cascades to regulate related but distinct forms of synaptic transmission across brain structures. We also observed an intriguing dichotomy between spontaneous versus evoked inhibition, as DAMGO elicited a greater suppression of eIPSCs than DPDPE. Several non-exclusive possibilities may account for these mixed effects. First, excitatory synapses can exhibit either spontaneous or evoked release [56, 60, 78], so spontaneous and evoked inhibition may preferentially sample inhibitory synapses that are predominantly regulated by DOR or MOR signaling, respectively. Second, an important caveat to our experimental preparation is that electrical stimulation elicits the release of a multitude of chemical messengers alongside GABA. It is therefore possible that electrical stimulation stimulates cell signaling that acts synergistically with DAMGO to suppress eIPSCs. For example, prior reports of endocannabinoid-MOR interactions at excitatory synapses suggest shared signaling pathways between CB1 receptors and MORs [7]. Finally, we cannot exclude that postsynaptic opioid signaling could contribute to reductions in eIPSC amplitude; however, this seems unlikely given the weak expression of MOR and DOR in L2/3 pyramidal cells.

### Somato-dendritic versus presynaptic regulation of GABA signaling by MOR and DOR

Nuances in the regulation of pre- and postsynaptic signaling can lead to important differences in how opioids regulate different circuits and behavior. We show that the balance of MOR or DOR signaling towards pre- or postsynaptic inhibition in the PL-PFC is shifted across IN types. The relationship between pre- and postsynaptic DOR signaling is linear, where the somatic response to DPDPE scales with the magnitude of presynaptic inhibition. In contrast, MOR activation had opposing effects on presynaptic versus postsynaptic signaling depending on the cell type. The most dramatic example of divergent pre- and postsynaptic function was observed in PV-INs, where MOR activation induced hyperpolarizing outward currents but did not alter PV-mediated oIPSCs. The converse was also observed, where CCK-IN oIPSCs were suppressed by MOR activation, but neither FS-CCK-INs or RS-CCK-INs exhibited DAMGO currents. One exciting explanation for these results is that MOR expression may be restricted to certain subcellular domains in PV-INs and CCK-INs. Future studies using advanced imaging techniques would be needed to rigorously test this hypothesis. Another interesting implication to these findings is that presynaptic MOR and DOR are less susceptible to desensitization than somato-dendritic receptors following prolonged receptor activation [19]. This is especially relevant in the context of opioid use disorder and related animal models that involve chronic opioid exposure. Receptor desensitization and internalization following repeated agonist treatment may differentially affect cells within which pre- and postsynaptic components of MOR signaling are skewed. For instance, PV-INs may be more susceptible to a loss of MOR signaling following prolonged receptor activation compared to cells with strong presynaptic signaling such as SST-INs.

One limitation of the CCK-IN optogenetic experiments is that we could not parse the effect of DAMGO and DPDPE on synaptic output from FS-CCK-INs versus RS-CCK-INs. The striking divergence in DOR sensitivity that we observed in whole-cell recordings from FS-CCK-INs and RS-CCK-INs suggests that there may also be differences in opioid regulation of their inhibitory output. Paired recordings from basket and pyramidal cells in the hippocampus have shown that GABA transmission from fast-spiking but not regular-spiking basket cells is inhibited by DAMGO [29], though the molecular identity of these cells is unclear. Future studies using more selective and/or combinatorial genetic approaches are needed to better define subtypes of CCK-INs.

Presynaptic inhibition of neurotransmitter release was measured by MOR- or DOR-induced suppression of oIPSC amplitude. Synaptic output from PV-INs and VIP-INs was inhibited by DOR or MOR signaling, respectively, while GABA release from SST-INs and CCK-INs can be reduced by both receptor systems. Opioid regulation of disinhibitory synapses may involve both presynaptic and postsynaptic signaling, providing a potential mechanism for differential DOR signaling at SST-PV and SST-PN synapses. Opioid-mediated plasticity of GABAergic transmission does not appear to be homologous across brain structures. Indeed, PV-IN and SST-IN synapses in the visual cortex are insensitive to MOR and DOR activation, respectively, while MOR and DOR both suppress hippocampal GABA signaling from these cells [15]. It is unclear whether other IN populations display similar patterns of opioid-associated plasticity across neo- and allocortex.

Despite the absence of pre- and postsynaptic MOR signaling in FSIs and RSNPs, a large proportion of MOR-EYFP+ cells were characterized as FSI or RSNP. Opioid receptors may alter neuronal excitability via AC or arrestin-mediated signaling, or kinases such as p38, ERK, PKC, and Src, leading to downstream changes in not reflected in the acute response to DAMGO or DPDPE. The large number of signaling pathways and downstream effectors coupled to these receptors necessitate additional pharmacological and electrophysiological assays to fully characterize opioid receptor function in these cell populations.

### Conclusions

GPCRs are targets for approximately one third of all FDA-approved medications for the treatment of psychiatric disorders [67]. MORs have emerged as effective targets for the treatment of substance use disorders-- especially OUD, though most opioid-based compounds are plagued by aversive side effects [23]. The DOR system has been heavily implicated in preclinical assessments of reward and negative affect [51, 71, 75], though the development of DOR-based therapeutics has been hindered by epileptic activity of first-generation DOR agonists [16]. The ongoing opioid epidemic has motivated recent advances in opioid receptor pharmacology which point towards molecular and cell type-specific drug development strategies as safer and more effective options.

We show that MOR and DOR control discrete aspects of inhibitory transmission through different signaling pathways, cellular functions, and circuit motifs in PFC, identifying several opportunities for potential cell- and synapse-restricted targets. *In vivo* characterization of how MOR and DOR guide IN function in health and disease will be the next step for translating these discoveries into next-generation psychiatric treatments.

## Author Contributions

RHC – Conceptualization, Investigation, Writing – Original Draft, Visualization; MEJ – Conceptualization, Investigation, Writing – Review & Editing, Supervision, Funding Acquisition.

## Acknowledgments

This work was supported by the National Institutes of Health [grant numbers R00AA027806, R01DA058702, and DP1DA060482], the Whitehall Foundation [grant number 2022-08-005], and the Brain and Behavior Research Foundation.

We thank members of the University of Pittsburgh Department of Psychiatry and Translational Neuroscience Program for stimulating discussions. We thank Gord Fishell and Yating Wang for providing the mDlx-FLEX-ChR2 plasmid. We thank Gregory Corder and Gregory Salimando for their technical advice and assistance regarding the MORp-EYFP virus.

## Competing Interests

The authors declare no competing interests.

## Notes

### Competing Interest Statement

The authors have declared no competing interest.

## References

1. Adzic, M., et al., Contribution of the opioid system to depression and to the therapeutic effects of classical antidepressants and ketamine. Life Sciences, 2023. 326: p. 121803.

2. Alexander, R.P. and K.J. Bender, Delta opioid receptors engage multiple signaling cascades to differentially modulate prefrontal GABA release with input and target specificity. bioRxiv, 2024.

3. Alexander, W.H. and T. Womelsdorf, Interactions of medial and lateral prefrontal cortex in hierarchical predictive coding. Frontiers in Computational Neuroscience, 2021. 15: p. 605271.

4. Arbabi, K., et al., Transcriptomic pathology of neocortical microcircuit cell types across psychiatric disorders. Molecular Psychiatry, 2024: p. 1–12.

5. Ariza, J., et al., The number of chandelier and basket cells are differentially decreased in prefrontal cortex in autism. Cerebral Cortex, 2018. 28(2): p. 411–420.

6. Artinian, J. and J.-C. Lacaille, Disinhibition in learning and memory circuits: New vistas for somatostatin interneurons and long-term synaptic plasticity. Brain research bulletin, 2018. 141: p. 20–26.

7. Atwood, B.K., D.A. Kupferschmidt, and D.M. Lovinger, Opioids induce dissociable forms of long-term depression of excitatory inputs to the dorsal striatum. Nature neuroscience, 2014. 17(4): p. 540–548.

8. Bacci, A., J.R. Huguenard, and D.A. Prince, Long-lasting self-inhibition of neocortical interneurons mediated by endocannabinoids. Nature, 2004. 431(7006): p. 312–316.

9. Bacci, A., J.R. Huguenard, and D.A. Prince, Modulation of neocortical interneurons: extrinsic influences and exercises in self-control. Trends in neurosciences, 2005. 28(11): p. 602–610.

10. Bandelow, B., et al., Borderline personality disorder: a dysregulation of the endogenous opioid system? Psychological review, 2010. 117(2): p. 623.

11. Bastien, G., et al., Effects of Buprenorphine/Naloxone and Methadone on Depressive Symptoms in People with Prescription Opioid Use Disorder: A Pragmatic Randomised Controlled Trial. The Canadian Journal of Psychiatry, 2023. 68(8): p. 572–585.

12. Birdsong, W.T., et al., Synapse-specific opioid modulation of thalamo-cortico-striatal circuits. Elife, 2019. 8: p. e45146.

13. Browne, C.A., M.L. Jacobson, and I. Lucki, Novel targets to treat depression: opioid-based therapeutics. Harvard review of psychiatry, 2020. 28(1): p. 40–59.

14. Bull, F.A., et al., Morphine activation of mu opioid receptors causes disinhibition of neurons in the ventral tegmental area mediated by β-arrestin2 and c-Src. Scientific reports, 2017. 7(1): p. 9969.

15. Caccavano, A.P., et al., Divergent opioid-mediated suppression of inhibition between hippocampus and neocortex across species and development. bioRxiv, 2024.

16. Chung, P.C.S., et al., Delta opioid receptors expressed in forebrain GABAergic neurons are responsible for SNC80-induced seizures. Behavioural brain research, 2015. 278: p. 429–434.

17. Clark, J.A., et al., [D-Pen2, D-Pen5] enkephalin (DPDPE): a δ-selective enkephalin with low affinity for μ1 opiate binding sites. European journal of pharmacology, 1986. 128(3): p. 303–304.

18. Cole, R.H., K. Moussawi, and M.E. Joffe, Opioid modulation of prefrontal cortex cells and circuits. Neuropharmacology, 2024: p. 109891.

19. Coutens, B. and S.L. Ingram, Key differences in regulation of opioid receptors localized to presynaptic terminals compared to somas: relevance for novel therapeutics. Neuropharmacology, 2023. 226: p. 109408.

20. Cummings, K.A. and R.L. Clem, Prefrontal somatostatin interneurons encode fear memory. Nature neuroscience, 2020. 23(1): p. 61–74.

21. Dienel, S.J. and D.A. Lewis, Alterations in cortical interneurons and cognitive function in schizophrenia. Neurobiology of disease, 2019. 131: p. 104208.

22. Dimidschstein, J., et al., A viral strategy for targeting and manipulating interneurons across vertebrate species. Nature neuroscience, 2016. 19(12): p. 1743–1749.

23. Edinoff, A.N., et al., Full opioid agonists and tramadol: pharmacological and clinical considerations. Anesthesiology and Pain Medicine, 2021. 11(4).

24. Fabian, C.B., et al., Parvalbumin interneuron mGlu5 receptors govern sex differences in prefrontal cortex physiology and binge drinking. Neuropsychopharmacology, 2024: p. 1–11.

25. Ferguson, B.R. and W.-J. Gao, PV interneurons: critical regulators of E/I balance for prefrontal cortex-dependent behavior and psychiatric disorders. Frontiers in neural circuits, 2018. 12: p. 37.

26. Fricker, L.D., et al., Five decades of research on opioid peptides: current knowledge and unanswered questions. Molecular pharmacology, 2020. 98(2): p. 96–108.

27. Fu, X., et al., Gq neuromodulation of BLA parvalbumin interneurons induces burst firing and mediates fear-associated network and behavioral state transition in mice. Nat Commun 13: 1290. 2022.

28. Ghosal, S., B.D. Hare, and R.S. Duman, Prefrontal cortex GABAergic deficits and circuit dysfunction in the pathophysiology and treatment of chronic stress and depression. Current opinion in behavioral sciences, 2017. 14: p. 1–8.

29. Glickfeld, L.L., B.V. Atallah, and M. Scanziani, Complementary modulation of somatic inhibition by opioids and cannabinoids. Journal of Neuroscience, 2008. 28(8): p. 1824–1832.

30. Gorelova, N., J.K. Seamans, and C.R. Yang, Mechanisms of dopamine activation of fast-spiking interneurons that exert inhibition in rat prefrontal cortex. Journal of neurophysiology, 2002. 88(6): p. 3150–3166.

31. Hashemi, E., et al., The number of parvalbumin-expressing interneurons is decreased in the prefrontal cortex in autism. Cerebral cortex, 2017. 27(3): p. 1931–1943.

32. He, M., et al., Strategies and tools for combinatorial targeting of GABAergic neurons in mouse cerebral cortex. Neuron, 2016. 91(6): p. 1228–1243.

33. Hippenmeyer, S., et al., A developmental switch in the response of DRG neurons to ETS transcription factor signaling. PLoS biology, 2005. 3(5): p. e159.

34. Horvath, P.M., et al., Spontaneous and evoked neurotransmission are partially segregated at inhibitory synapses. Elife, 2020. 9: p. e52852.

35. Hou, G., et al., Opioid receptors modulate firing and synaptic transmission in the paraventricular nucleus of the thalamus. Journal of Neuroscience, 2023. 43(15): p. 2682–2695.

36. Jiang, C., et al., Morphine coordinates SST and PV interneurons in the prelimbic cortex to disinhibit pyramidal neurons and enhance reward. Molecular psychiatry, 2021. 26(4): p. 1178–1193.

37. Joffe, M.E., et al., Acute restraint stress redirects prefrontal cortex circuit function through mGlu5 receptor plasticity on somatostatin-expressing interneurons. Neuron, 2022. 110(6): p. 1068–1083. e5.

38. Joffe, M.E., D.G. Winder, and P.J. Conn, Contrasting sex-dependent adaptations to synaptic physiology and membrane properties of prefrontal cortex interneuron subtypes in a mouse model of binge drinking. Neuropharmacology, 2020. 178: p. 108126.

39. Kaar, S.J., et al., Pre-frontal parvalbumin interneurons in schizophrenia: a meta-analysis of post-mortem studies. Journal of Neural Transmission, 2019. 126: p. 1637–1651.

40. Kaiser, T., et al., Transgenic labeling of parvalbumin-expressing neurons with tdTomato. Neuroscience, 2016. 321: p. 236–245.

41. Kamalova, A., et al., CCK+ Interneurons Contribute to Thalamus-Evoked Feed-Forward Inhibition in the Prelimbic Prefrontal Cortex. Journal of Neuroscience, 2024. 44(23).

42. Kawaguchi, Y., Selective cholinergic modulation of cortical GABAergic cell subtypes. Journal of neurophysiology, 1997. 78(3): p. 1743–1747.

43. Kawaguchi, Y. and T. Shindou, Noradrenergic excitation and inhibition of GABAergic cell types in rat frontal cortex. Journal of Neuroscience, 1998. 18(17): p. 6963–6976.

44. Kullander, K. and L. Topolnik, Cortical disinhibitory circuits: cell types, connectivity and function. Trends in Neurosciences, 2021. 44(8): p. 643–657.

45. Lau, B.K., et al., Mu-opioids suppress GABAergic synaptic transmission onto orbitofrontal cortex pyramidal neurons with subregional selectivity. Journal of Neuroscience, 2020. 40(31): p. 5894–5907.

46. Law, P.-Y., Y.H. Wong, and H.H. Loh, Molecular mechanisms and regulation of opioid receptor signaling. Annual review of pharmacology and toxicology, 2000. 40(1): p. 389–430.

47. Lee, S., et al., A disinhibitory circuit mediates motor integration in the somatosensory cortex. Nature neuroscience, 2013. 16(11): p. 1662–1670.

48. Lewis, D.A., et al., Cortical parvalbumin interneurons and cognitive dysfunction in schizophrenia. Trends in neurosciences, 2012. 35(1): p. 57–67.

49. Lupica, C.R., Delta and mu enkephalins inhibit spontaneous GABA-mediated IPSCs via a cyclic AMP-independent mechanism in the rat hippocampus. Journal of Neuroscience, 1995. 15(1): p. 737–749.

50. Lupica, C.R., W.R. Proctor, and T.V. Dunwiddie, Dissociation of μ and δ opioid receptor-mediated reductions in evoked and spontaneous synaptic inhibition in the rat hippocampus in vitro. Brain research, 1992. 593(2): p. 226–238.

51. Lutz, P.-E. and B.L. Kieffer, Opioid receptors: distinct roles in mood disorders. Trends in neurosciences, 2013. 36(3): p. 195–206.

52. Ma, X., et al., In vivo photopharmacology with a caged mu opioid receptor agonist drives rapid changes in behavior. Nature methods, 2023. 20(5): p. 682–685.

53. Madisen, L., et al., A robust and high-throughput Cre reporting and characterization system for the whole mouse brain. Nature neuroscience, 2010. 13(1): p. 133–140.

54. Maksymetz, J., et al., mGlu1 potentiation enhances prelimbic somatostatin interneuron activity to rescue schizophrenia-like physiological and cognitive deficits. Cell reports, 2021. 37(5).

55. Margolis, E.B. and H.L. Fields, Mu opioid receptor actions in the lateral habenula. PloS one, 2016. 11(7): p. e0159097.

56. Melom, J.E., et al., Spontaneous and evoked release are independently regulated at individual active zones. Journal of Neuroscience, 2013. 33(44): p. 17253–17263.

57. New, A.S. and B. Stanley, An opioid deficit in borderline personality disorder: self-cutting, substance abuse, and social dysfunction. 2010, Am Psychiatric Assoc. p. 882–885.

58. Oh, D.H., et al., Neuropathological abnormalities of astrocytes, GABAergic neurons, and pyramidal neurons in the dorsolateral prefrontal cortices of patients with major depressive disorder. European Neuropsychopharmacology, 2012. 22(5): p. 330–338.

59. Patton, M.H., et al., Ethanol disinhibits dorsolateral striatal medium spiny neurons through activation of a presynaptic delta opioid receptor. Neuropsychopharmacology, 2016. 41(7): p. 1831–1840.

60. Peled, E.S., Z.L. Newman, and E.Y. Isacoff, Evoked and spontaneous transmission favored by distinct sets of synapses. Current Biology, 2014. 24(5): p. 484–493.

61. Pi, H.-J., et al., Cortical interneurons that specialize in disinhibitory control. Nature, 2013. 503(7477): p. 521–524.

62. Puryear, C.B., et al., Opioid receptor modulation of neural circuits in depression: What can be learned from preclinical data? Neuroscience & Biobehavioral Reviews, 2020. 108: p. 658–678.

63. Raveh, A. and E. Reuveny, Mechanisms of Short-Term Desensitization of GIRK Channel Activity. Biophysical Journal, 2009. 96(3): p. 463a.

64. Reeves, K.C., et al., Opioid receptor-mediated regulation of neurotransmission in the brain. Frontiers in Molecular Neuroscience, 2022. 15: p. 919773.

65. Salimando, G.J., et al., Human OPRM1 and murine Oprm1 promoter driven viral constructs for genetic access to μ-opioidergic cell types. Nature Communications, 2023. 14(1): p. 5632.

66. Schmauss, C. and H.M. Emrich, Dopamine and the action of opiates: A reevaluation of the dopamine hypothesis of schizophrenia with special consideration of the role of endogenous opioids in the pathogenesis of schizophrenia. Biological psychiatry, 1985. 20(11): p. 1211–1231.

67. Schmitz, G.P. and B.L. Roth, G protein-coupled receptors as targets for transformative neuropsychiatric therapeutics. American Journal of Physiology-Cell Physiology, 2023. 325(1): p. C17–C28.

68. Seamans, J.K., et al., Bidirectional dopamine modulation of GABAergic inhibition in prefrontal cortical pyramidal neurons. Journal of Neuroscience, 2001. 21(10): p. 3628–3638.

69. Smith, S.J., et al., Single-cell transcriptomic evidence for dense intracortical neuropeptide networks. elife, 2019. 8: p. e47889.

70. Stein, C., Opioid Receptors. Annu Rev Med, 2016. 67: p. 433–51.

71. Suzuki, T., et al., Blockade of δ-opioid receptors prevents morphine-induced place preference in mice. The Japanese Journal of Pharmacology, 1994. 66(1): p. 131–137.

72. T Lamberts, J. and J. R Traynor, Opioid receptor interacting proteins and the control of opioid signaling. Current pharmaceutical design, 2013. 19(42): p. 7333–7347.

73. Taki, K., T. Kaneko, and N. Mizuno, A group of cortical interneurons expressing μ-opioid receptor-like immunoreactivity: a double immunofluorescence study in the rat cerebral cortex. Neuroscience, 2000. 98(2): p. 221–231.

74. Taniguchi, H., et al., A resource of Cre driver lines for genetic targeting of GABAergic neurons in cerebral cortex. Neuron, 2011. 71(6): p. 995–1013.

75. Torregrossa, M.M., et al., Peptidic delta opioid receptor agonists produce antidepressant-like effects in the forced swim test and regulate BDNF mRNA expression in rats. Brain research, 2006. 1069(1): p. 172–181.

76. Tremblay, R., S. Lee, and B. Rudy, GABAergic interneurons in the neocortex: from cellular properties to circuits. Neuron, 2016. 91(2): p. 260–292.

77. Trigo, J.M., et al., The endogenous opioid system: a common substrate in drug addiction. Drug and alcohol dependence, 2010. 108(3): p. 183–194.

78. Walter, A.M., V. Haucke, and S.J. Sigrist, Neurotransmission: spontaneous and evoked release filing for divorce. Current Biology, 2014. 24(5): p. R192–R194.

79. Xu, H., et al., A disinhibitory microcircuit mediates conditioned social fear in the prefrontal cortex. Neuron, 2019. 102(3): p. 668–682. e5.

80. Zamfir, M., et al., Distinct and sex-specific expression of mu opioid receptors in anterior cingulate and somatosensory S1 cortical areas. Pain, 2023. 164(4): p. 703.

81. Zaratin, P.F., et al., Modification of nociception and morphine tolerance by the selective opiate receptor-like orphan receptor antagonist (–)-cis-1-methyl-7-[[4-(2, 6-dichlorophenyl) piperidin-1-yl] methyl]-6, 7, 8, 9-tetrahydro-5H-benzocyclohepten-5-ol (SB-612111). Journal of Pharmacology and Experimental Therapeutics, 2004. 308(2): p. 454–461.

